# *Gexplora* – user interface that highlights and explores the density of genomic elements along a chromosomal sequence

**DOI:** 10.1101/2020.04.04.025379

**Authors:** Thomas Nussbaumer, Olivia Debnath, Parviz Heidari

## Abstract

The density of genomic elements such as genes or transposable elements along its consecutive sequence can provide an overview of a genomic sequence while in the detailed analysis of candidate genes it may depict enriched chromosomal hotspots harbouring genes that explain a certain trait. The herein presented python-based graphical user interface *Gexplora* allows to obtain more information about a genome by considering sequence-intrinsic information from external databases such as Ensembl, OMA and STRING database using REST API calls to retrieve sequence-intrinsic information, protein-protein datasets and orthologous groups. Gexplora is available under https://github.com/nthomasCUBE/Gexplora.

## Introduction

In the last years we observed a steady increase in the number of sequenced genomes granted by improvements and release of sequencing technologies, assembly strategies that consider shorter and longer reads, hybrid assemblies ^12^ and optical maps to merge scaffolds ^1^. Other technical aspects include the improvements of computational tools for read assembly ^11^ and strategies to filter reads prior to analysis ^13^ as well as approaches to perform gap-filling ^14^. This along with reductions in sequencing costs and the requirement to focus more intensively on the biology of a novel genome - as sequencing has become feasible for many research groups - has led to a shift from single genome focussed analysis towards analysis of meta-genomes and pan-genomes ^2^ and thereby allowed to draw a broader picture that explains genomic or transcriptional differences between cultivars, wild types or subpopulations.

A relevant example for a large and challenging genome is bread wheat (*Triticum aestivum*) where the genome size exceeds the human genome by fivefold. Thereby, in the last years, research consortia obtained great success in determining the draft genome and later its near-complete linear genomic template ^3^. In case that a genome was successfully assembled into chromosomes, underlying datasets are typically deposit in central repositories such as Ensembl ^4^ or Orthologous Matrix Database ^5^ which enable a collection of genomes or gene annotations from multiple species while in the field of microbes frequently used resources include eggNOG ^9^ and NCBI Microbial Genomes (https://www.ncbi.nlm.nih.gov/genome/microbes/).

The analysis of genome structure and gene content is key to assess the completeness or relatedness of a genome of interest and there exist plenty tools to compare the density of genes overall, e.g. the decay of orthologous genes or to derive syntenic blocks that can be then visualized with tools such as Circos ^6^ or CrowsNest ^7^, among many other supportive tools which help to understand the evolution of the species when facing introgression events or when related genomes are compared to a common ancestor.

In order to analyse the density of candidate and gene density in crops, in a previous study, we have implemented *chromoWIZ* ^10^ as a web service to visualize gene densities along their genomes with a focus on four selected plant genomes namely *Brachypodium distachyon, Hordeum vulgare, Triticium aestivum* and *Oryza sativa*. In the current version, we provide more features with the possibility to consider any genome where a gene transfer formatted (GTF) file is available as well as selected interfaces to public databases such as OMA, STRING database and Ensembl that account for the most commonly used web resources for obtaining and analyzing genomic data.

In the last two decades’ genomics and genetic resources were mainly accessible by the institutes who coordinated the study including a FTP repository and ideally web forms to query the data. However, a comparison between different datasets was often difficult due to slightly different data standards. Meanwhile, a more efficient and common way to provide genomic datasets is to share and integrate data files between tools on the basis of REpresentational State Transfer API (REST API) ^8^ calls that often provide JSON-formatted methods (‘endpoints’) that allow to request (*‘GET’*), update (*‘PUT’*) or send data (*‘POST’*). Many resources provide such endpoints which can be then integrated into customized tools or scripts. To demonstrate this for the analysis of gene distributions, we used selected endpoints from the three resources: *Gexplora* integrates endpoints from Ensembl in order to obtain sequence-based information while we used OMA to retrieve more information about orthologous proteins and string-DB to determine associated publications and protein-protein interactions.

## Material and Methods

### Calculation of the density of genetic elements

The distribution of each element type (*e.g.* gene or mRNA) can be selected after the GTF (gene transfer format) file has been uploaded into *Gexplora*. The density of an element type is shown per bin which represents one visualization unit and indicates 1/200 of the chromosome length by considering the largest genome. Bins, with the highest amount of elements are shown in *red* while the low amounts of elements are shown in *blue. Gexplora* depicts the first five chromosomes. For genomes with more chromosomes, the next five chromosomes can be shown by activating ‘forward’ or ‘reverse’ buttons. The maximum density can be set by the user in the top panel of the tool. If the density of the candidate genes should be shown, this is possible via the option ‘*Find gene*’ allowing to upload the candidate genes. The graphical user interface *Gexplora* is implemented using the package *‘Tkinter’* package that is available in the default python installation while several additional python packages including *numpy, requests* and *json* to parse the response objects from three resources. For the export of the densities per chromosomes we make use of the Python package *XlsxWriter* ^16^. This can be done with the executable *‘pip.exe’* in Windows that is located in the ‘Scripts’ package of the Python installation directory. For the graphical user interface the Python version 3.7 was used along with Windows as the operating system.

### Obtaining data through REST API calls

We have used three external databases that provide REST APIs: Ensembl, STRING database and OMA Orthology Database (Figure 1). For Ensembl, we allow to obtain sequences for individual genes (entrypoint: /family/member/id/<gene>). From OMA we obtain the gene family that contains the candidate gene (method: *orthologs*) while STRING database allows to retrieve publications that are associated with the gene (abstractsList).

**Figure 1.**
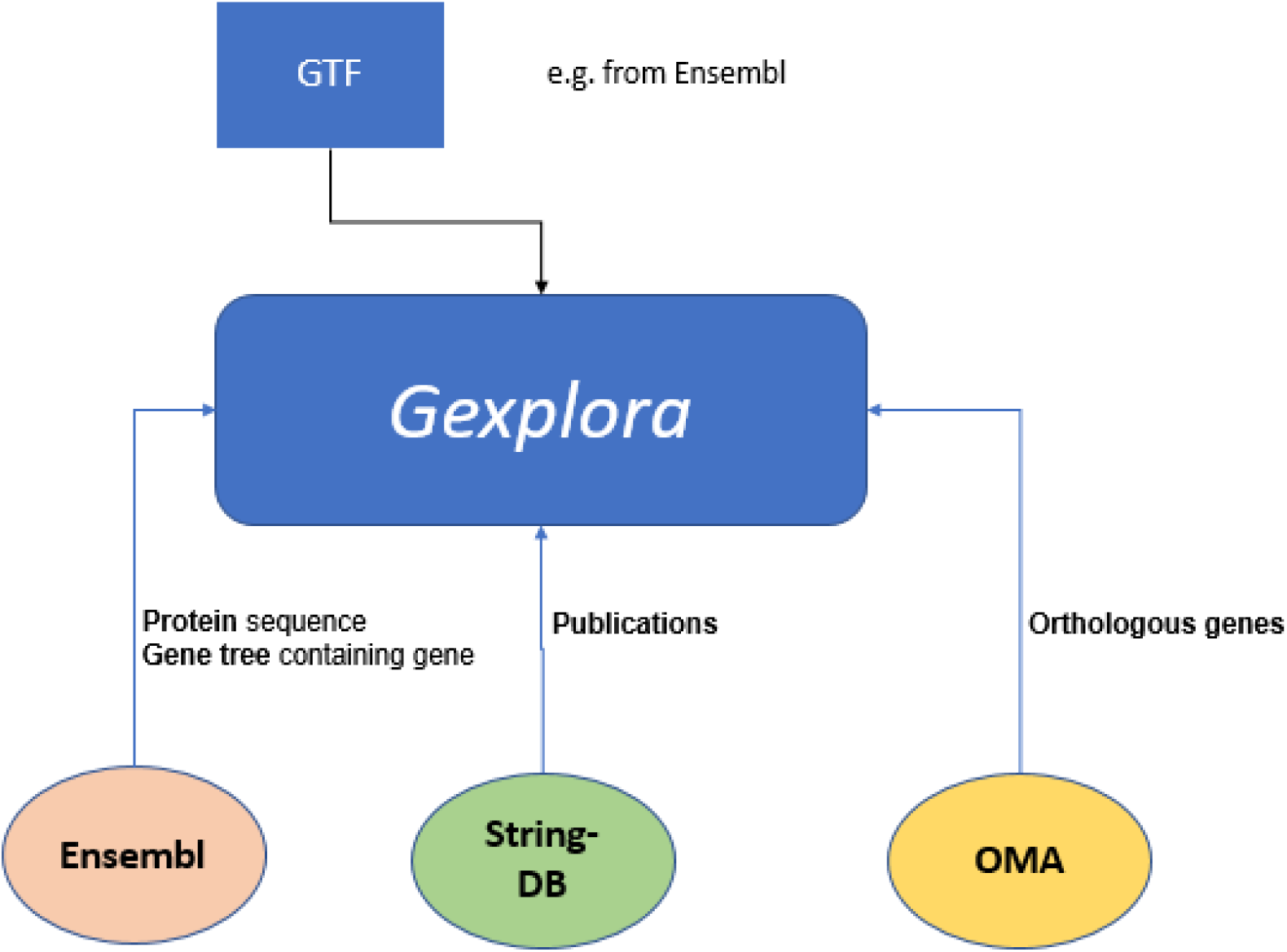
*Gexplora* workflow. The workflow illustrates the functionality of *Gexplora* where a user can upload a gene transfer file (GTF) as input and where interfaces to Ensembl, STRING database and OMA exist.

### Extraction of the data described in the show cases

We have downloaded the GTF file from Ensembl (ftp://ftp.ensemblgenomes.org/pub/release-46/plants/gff3/brachypodium_distachyon).

## Results

In our last study that we have published several years ago ^10^, we have described the web-server *chromoWIZ* to display gene distributions along four selected and agriculturally important or model plants namely *Brachypodium distachyon, Triticum aestivum, Hordeum vulgare* and *Oryza sativa*. As the analysis of the density of element types might be also of interest for many other genomes or genomes where annotation files are regularly updated, we provide most of the features also in *Gexplora* within a graphical user interface with the option to change the annotation file (see Table 1, Figure 1). These annotation files can be downloaded from Ensembl. Apart from the basic functions to visualize the element density, we offer export functions (e.g. statistics of the amount of elements per chromosomes) and interfaces to three commonly used web repositories Ensembl, OMA and STRING database.

**Table 1.** Overview of the various tools and when compared to the previous version of the tool.

| Aspect | *chromoWIZ* | *Gexplora* |
| --- | --- | --- |
| Species to analyse | *Brachypodium distachyon, Triticum aestivum, Hordeum vulgare, Oryza sativa* | any genome with a GTF (gene transfer format) files of genetic elements |
| Features | Mapping of genes to chromosomes<br>Enrichment analysis for all chromosomes whether contain more genes than expected | Mapping of genes to chromosomes<br>(Enrichment analysis whether genes are enriched in certain chromosomes) |
| Integration to Ensembl | *n.a.* | Retrieval of sequences and gene trees of one or several candidate genes with Ensembl REST API<br>Orthologous genes with OMA<br>Pubmed Identifiers referencing a candidate gene with string-DB |

While OMA allows to analyse orthologous proteins in comparison to other species; STRING database allows to analyse protein-protein interactions as well as provides literature entries that reference a gene among many other features. The REST API of Ensembl we have used to retrieve a gene tree for a candidate gene. This can be helpful, when a researcher wants to obtain the orthologous genes, e.g. compare whether there are differences in the copy numbers compared to other closely or distantly related species. This is provided by two endpoints as provided by Ensembl using REST API calls.

One of the big challenges in using different genomic web repositories is that gene identifiers change frequently. Otherwise, genomic repositories have their own gene identifiers. For instance, OMA uses its own gene identifiers, while STRING database uses public gene identifiers and Ensembl uses its own identifiers as well. To allow comparison and integration of the tools, mapping of identifiers is crucial. Fortunately, all these resources also provide the gene names, which was a central feature in *Gexplora* to allow comparison where a user can enable to also search using the gene name.

### Use case - Analysing the model plant *Brachypodium distachyon*

To benchmark our tool, we used the annotation of *Brachypodium distachyon* that contains five chromosomes. After the annotation file was uploaded, various element types appear (e.g. mRNA, tRNA) and can be selected. Upon selection, this triggers redrawing the overall density of the selected element type (e.g. ‘*mRNA*’ in Figure 1). We also offer an upload of candidate genes where after upload, the tool depicts the distribution of candidate genes along the chromosome using the file upload that is provided in ‘*Find genes*’ (Figure 2). For particular genes, we allow to obtain the coding sequences from Ensembl as well as a gene tree that contains orthologous and homologous genes to closely related species (Figure 2BC). By considering the STRING database, we are able to search for literature referencing these genes where candidate genes are described while the OMA interface allows to obtain all orthologous proteins.

**Figure 2.**
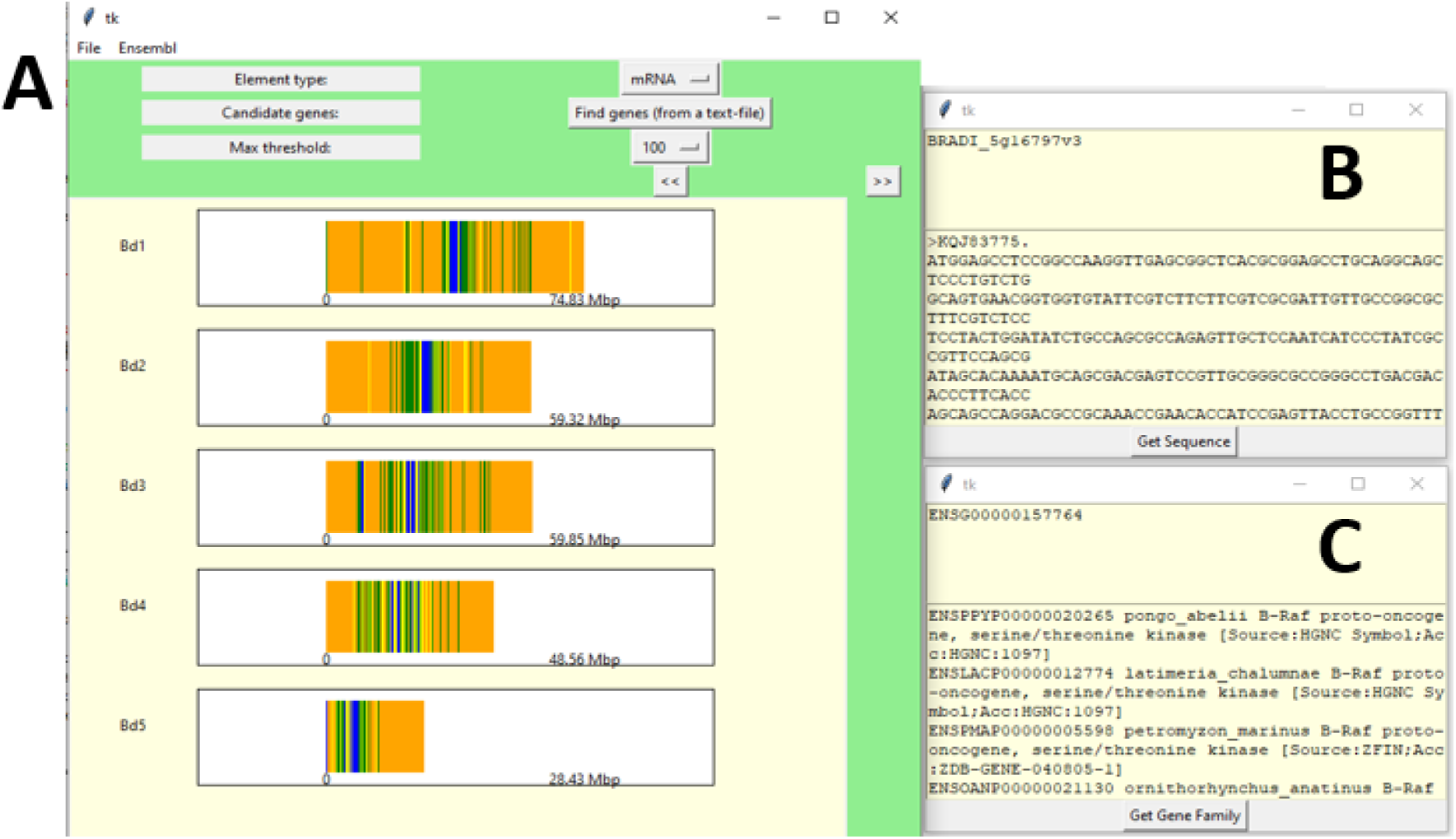
Gexplora. *Gexplora* takes an annotation file in GTF (gene transfer format) file as input (File -> Load). Main front end is shown in subfigure A, while subfigure B shows the coding sequence of a candidate gene as a result from an Ensembl search while subfigure C shows all genes that share homology to the given human candidate genes using Ensembl.

## Conclusion

*Gexplora* as a user-friendly interface tool provides a global overview of genomic elements of candidate genes using Ensembl, STRING database, and OMA and allows to display the density of genetic elements but also various features to obtain information of orthologous genes and sequences from three databases. As these databases offer many more methods and because we provide the source code of the tool, the interfaces can be adapted easily.

## Availability

*Gexplora* is available under: https://github.com/nthomasCUBE/Gexplora.

